# Sensory neurons control heritable adaptation to stress through germline reprogramming

**DOI:** 10.1101/406033

**Authors:** Giusy Zuco, Vikas Kache, Pedro Robles, Jyotiska Chaudhuri, Beth Hill, Christine Bateson, Andre Pires-daSilva

**Affiliations:** Department of Biology, University of Texas at Arlington, Arlington, TX, 76019, USA; School of Life Sciences University of Warwick, Coventry, CV4 7AL, UK (A.P.-d.S.)

**Author notes:** Corresponding author: Correspondence and requests for materials should be addressed to A.P.-d.S. Contributed equally (Giusy Zuco, Vikas Kache, Pedro Robles).

## Abstract

Maternal neuronal signaling has been reported to program adaptive changes in offspring physiology in diverse organisms [1, 2]. However, the mechanisms for the inheritance of adaptive maternal effects through the germline are largely unknown. In the nematode *Auanema freiburgensis*, stress-resistance and sex of the offspring depend on environmental cues experienced by the mother. Maternal sensing of high population densities results in the production of stress-resistant larvae (dauers) that develop into hermaphrodites. Ablation of the maternal ASH chemosensory neurons results only in non-dauer offspring that develop into males or females. High population densities correlate with changes in the methylation status of H3K4 and H3K9 in the maternal germline. Inhibition of JMJD histone demethylases prevents mothers from producing dauers and hermaphrodite offspring in high-density conditions. Our results demonstrate a case of soma-to-germline transmission of environmental information that influences the phenotype of the following generation through changes in histone modifications of the maternal germline.

**Highlights:**

- High population density leads to the production of hermaphrodite offspring.
- The ASH neuron in the hermaphrodite mother senses population density.
- Histone modifications in the maternal germline correlate with the sex of offspring.
- Inhibition of histone demethylases results in female offspring in all conditions.

---

Mechanisms for passing information about the maternal environment to the offspring evolved in several organisms. They allow mothers to match the phenotype of their offspring to changes in local environment, increasing their fitness. For example, mothers of the crustacean *Daphnia cucullata* and of some rotifer species generate predator-resistant offspring when sensing a predator cue [3, 4], seasonal changes sensed by some insect mothers result in stress-resistant offspring [5], and high population densities experienced by the red squirrel mother results in faster growing offspring that are more likely to acquire a territory and survive their first winter [2]. However, the mechanisms involved in the transmission of the environmental information to the following generation are largely unknown.

In populations of *Auanema* nematodes, three sexes coexist: XX hermaphrodites, XX females and XO males [6]. The male sex is chromosomally determined [7, 8], whereas the mechanism of hermaphrodite versus female sex determination is largely unknown. A crucial factor in the development of hermaphrodites in this nematode genus is the passage through the stress-resistant dauer stage [6, 9, 10], which has morphological and behavioral adaptations for dispersal. In *A*. *freiburgensis*, XX larvae that pass through the dauer stage always become hermaphrodites (N= 96), whereas XX non-dauer larvae develop into females (N= 93).

In *A*. *freiburgensis*, the environment of the mother determines the sexual fate of the XX offspring: hermaphrodite individuals kept in isolation produce mostly female offspring (99.4% ± 0.6% SE, N= 149 F1 offspring from 10 mothers), whereas hermaphrodites exposed to high nematode density conditions produce mostly hermaphrodite offspring (86.7% ± 2.4% SE, N= 199 F1 offspring from 10 mothers). In these experiments, high-density conditions were induced by incubating nematodes with conditioned medium (CM) of high-density *A*. *freiburgensis* liquid cultures (see Methods). Importantly, only the parental generation was exposed to the conditioned medium. Thus, these results suggest that dauer formation in *A*. *freiburgensis* is induced across a generation, instead of within the same generation as in *Caenorhabditis elegans* [11] (Figure 1A). The induction of dauers through the hermaphrodite mother is limited to one generation: F1 hermaphrodites derived from mothers in (+) CM plates produce mostly female offspring (99.6% ± 0.3% SE, N= 470 F2 offspring from 10 F1s).

**Figure 1.**
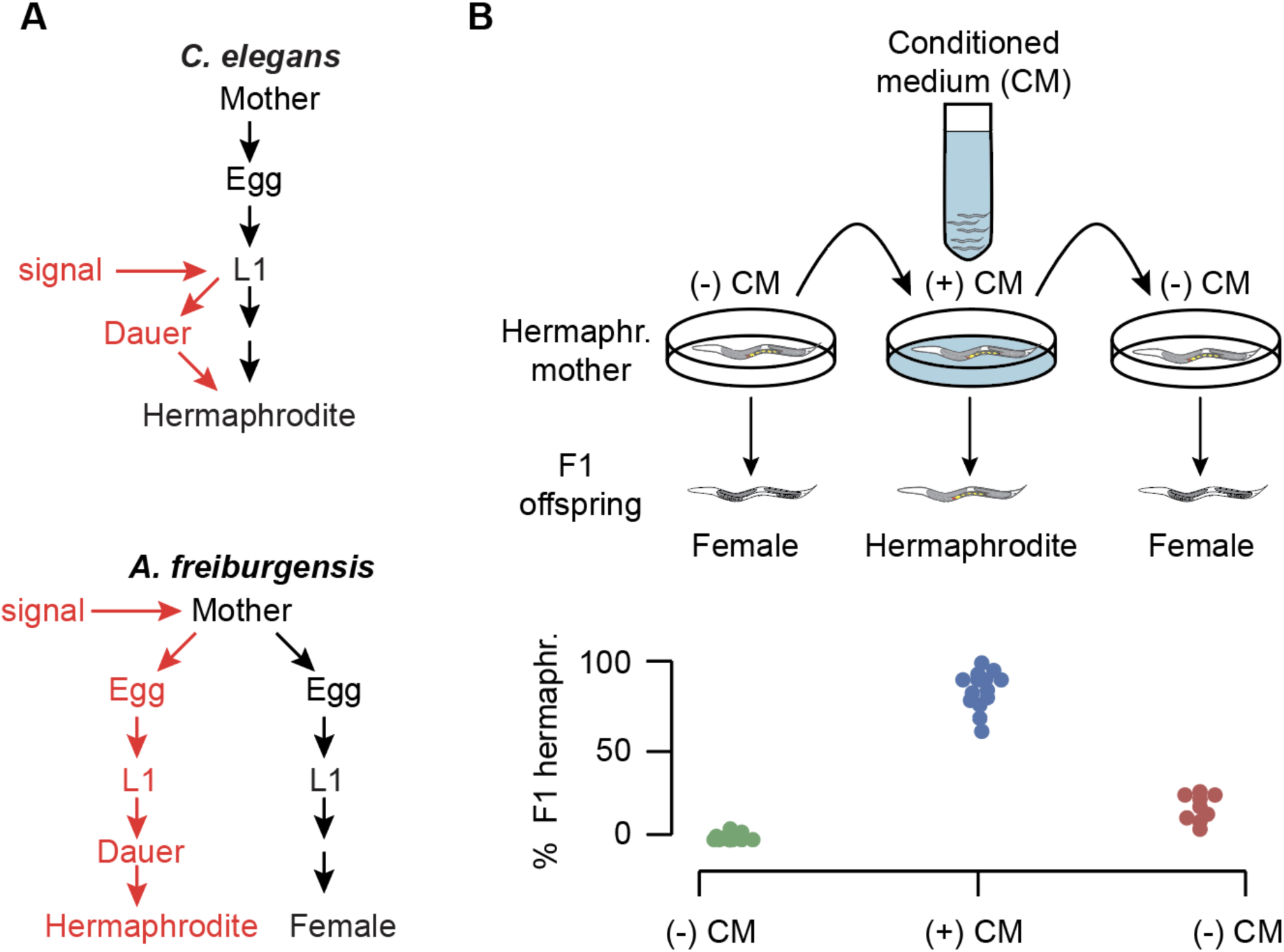
Dauer and hermaphrodite development are induced across generations in *A*. *freiburgensis*. **A.** In *C*. *elegans*, the L1 larvae respond to environmental signals to facultatively form stress-resistant dauers. *A*. *freiburgensis*, it is the mother and not the L1s that respond to environmental signals. *A*. *freiburgensis* dauers larvae obligatorily develop into hermaphrodite adults. **B.** In the experimental setup (top), the same individual mother hermaphrodite was transferred every 24 hours to a new environmental condition. Initially, it was placed in a plate without conditioned medium (-) CM, followed by the transfer to a (+) CM plate and then to a new (-) CM plate. The sex of the offspring was then assessed. Mothers (N= 14) kept in a (-) CM plate produced 1.7% of hermaphrodites (N total offspring= 386). When the same mothers (N= 14) were transferred to a (+) CM plate, they generated a mean of 83% of F1 hermaphrodites (N total offspring= 415). After transferring back to a new (-) CM plate, mothers (N= 10, 4 died) produced 17% F1 hermaphrodites (N total offspring= 364).

To test if *A*. *freiburgensis* L1 larvae can also respond to crowding conditions, similar to *C*. *elegans* L1 larvae, eggs derived from mothers cultured in isolation were left to hatch and undergo larval development in (+) CM plates until adulthood. 95.7% (N= 161) of these L1s developed into females, indicating that larvae do not respond to crowding conditions.

To determine if the CM affects dauer formation and sex determination in a reversible manner, individual mothers were tested in different conditions throughout their lives (Figure 1B). Eggs laid by hermaphrodites on (-) CM plates developed into females. When the same adult individuals were transferred to (+) CM plates, the offspring were mostly hermaphrodite. After rinsing the same mothers with M9 buffer and placing them onto a new (-) CM plate for about 24 hours, most offspring developed into females again. These results indicate that a hermaphrodite mother can reversibly respond to the environmental conditions.

When cultured for 6 hours, a minimum density of 16 adult hermaphrodites per cm^2^ is sufficient for the induction of 100% (N= 295) of hermaphrodite offspring. In densities below 10 individuals/cm^2^, the hermaphrodite mothers produce practically only female offspring (10 individuals/cm^2^, 100% F1 female, N= 78; 6 individuals/cm^2^, 98.5% F1 female, N= 66). At an intermediate density (13 individuals/cm^2^), hermaphrodites produce 19% (N= 126) of hermaphrodite offspring.

In *C*. *elegans*, other environmental stresses, such as incubation at high temperature (25 °C) or lack of food, can induce L1s to develop into dauers [11]. However, a 24-hour exposure of *A*. *freiburgensis* hermaphrodite mothers to high temperature or starvation resulted in mostly (97%) non-dauer larvae offspring that developed into female adults, for both conditions (N= 166 F1s from mothers at 25 °C and N= 146 F1s from starving mothers).

Nematodes have bilateral pairs of sensory organs in the head called amphids. In *C*. *elegans*, some of these amphid neurons are necessary to sense the environment and regulate dauer development [12]. To test if this was also the case for *A*. *freiburgensis* adults, we first identified each amphid. Amphid neurons have open sensory endings and thus take up lipophilic dyes such as DiI from the environment, allowing the visualization of their cell bodies [13]. Based on their relative position and by using *C*. *elegans* and other nematodes as reference [14], we identified ASK, ADL and ASH as the amphids that stain with DiI in *A*. *freiburgensis* (Figure 2A). We systematically ablated each pair type by using a laser microbeam. Laser ablation of the two ASH neurons in the mother hermaphrodite kept in a (+) CM plate was sufficient to prevent the production of dauer and hermaphrodite offspring (Figure 2B).

**Figure 2.**
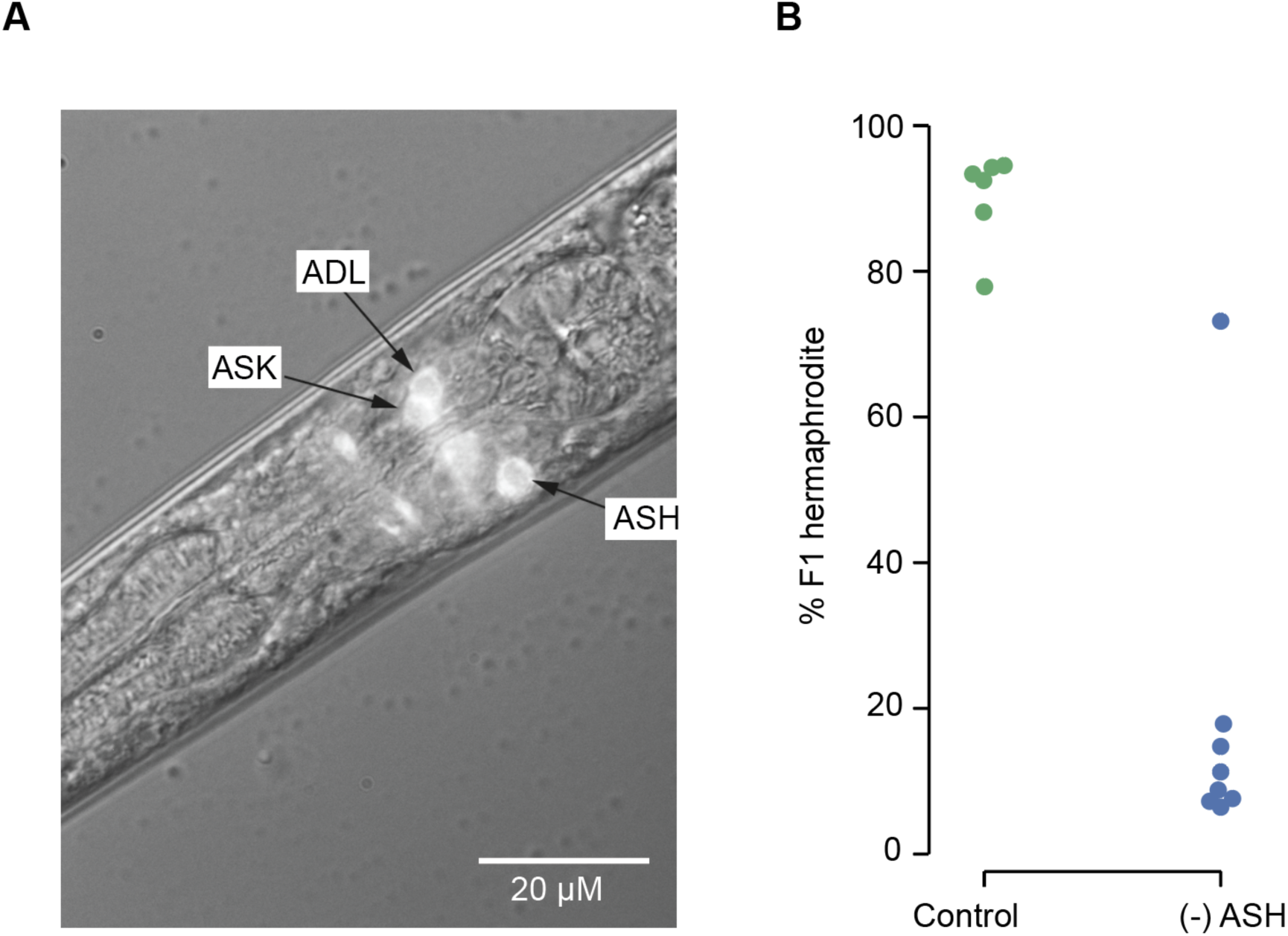
Killing of the neuronal pair ASH by laser ablation prevents the mother from producing hermaphrodite offspring. **A.** Amphid neurons stained with DiI were identified by the relative location of their cell bodies. **B.** When cultured in (+) CM plates, mock-ablated hermaphrodites (N= 6) generate mostly hermaphrodite offspring (90%, N total offspring= 553). In contrast, hermaphrodites in which both ASH neurons were ablated (N= 8) and kept in (+) CM plates, produced fewer hermaphrodite offspring (18%, N total offspring= 664). The outlier that produced a high proportion of hermaphrodite F1s is likely to be an animal in which only one of the ASH neurons was successfully ablated.

## Changes in histone modifications in the germline correlate with exposure to crowding conditions

To test if exposure of *A*. *freiburgensis* hermaphrodites to CM changes the epigenetic status of the chromatin and thus transcriptional activity in the germline, we performed antibody staining to detect histone modifications. We examined histone modifications that result in the methylation of lysine (K) residues of the histones 3 (H3) and 4 (H4) [15]. Similarly to *C*. *elegans* [16], the distal tip of the *A*. *freiburgensis* germline is mitotically active, whereas the remaining cells undergo meiosis. In gonads isolated from hermaphrodites grown in (-) CM plates, most animals have uniform levels of H3K4me3 (100%, N= 44 gonads) and H3K9me3 (73%, N= 67) throughout the germline (Figure 3). In gonads derived from animals in (+) CM plates, we observed clustered cells with high levels of H3K4me3 (76.3%, N= 59) and low levels of H3K9me3 (81%, N= 91) in the mitotic region (Figure 3).

**Figure 3.**
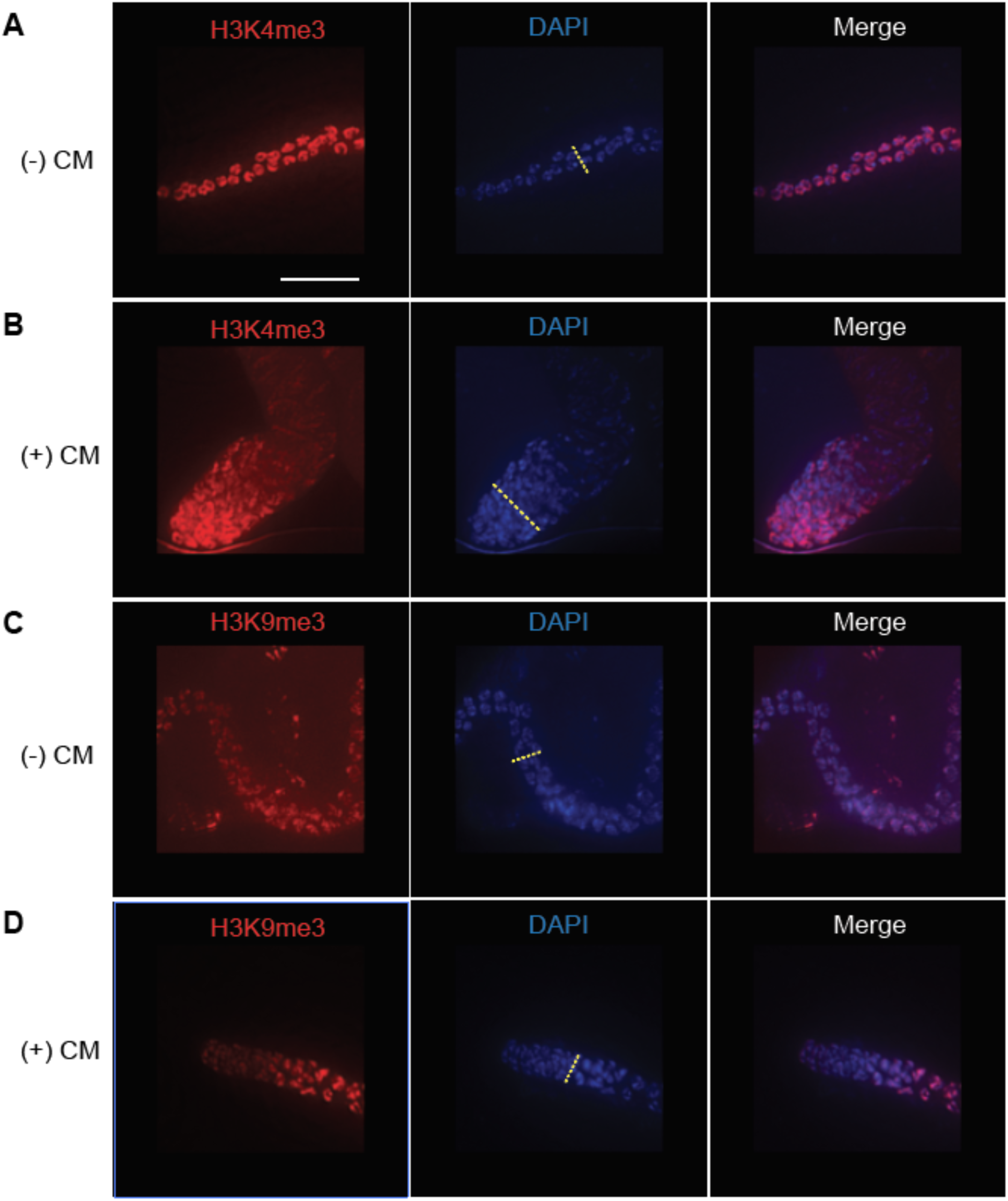
Histone methylation patterns change when hermaphrodite mothers are in (+) CM plates. When cultured in the (-) CM plates, hermaphrodites have homogeneous levels of H3K4me3 (**A**) and H3K9me3 (**B**) along the germline. When cultured in (+) CM plates, there is a higher level of H3K4me3 (**C**) and lower levels of H3K9me3 (**D**) in the tip of the gonad. The yellow dotted line in the pictures with DAPI staining represents the border between the mitotic and meiotic part of the germline. The mitotic and meiotic part of the gonad is to the left and the right of the yellow dotted line, respectively. Scale bar= 15 μm.

To test if those histone modifications have functional relevance, we used an inhibitor of Jumonji demethylases involved in H3K9me3 demethylation [17, 18]. The prediction is that treatment of animals with the KDM4/JMJD-2 inhibitor IOX-1 [18] would increase the rate of H3K9me3 in animals exposed to CM. This would reflect in lower number of hermaphrodites produced when mothers are exposed to CM. Accordingly, we found that most mothers treated with CM and 1 μM IOX-1 have higher levels of H3K9me3 (57.1%, N= 7 gonads) and lower levels of H3K4me3 (64.7%, N= 17 gonads) in the mitotic region (Supplementary Figure S1). They also produced mostly females (68%, N= 341 sexed F1s). In control experiments, in which hermaphrodites were treated only with CM, they produced mostly hermaphrodites (82%, N= 45 sexed F1s).

There are many examples of non-genetic mechanisms that induce phenotypes across generations [3, 19-21]. However, it is often difficult to determine whether offspring phenotypes are a passive consequence of resource availability to the parental generation or the result of an adaptive response across generations [20]. Furthermore, the mechanisms of transducing an environmental signal from the parental generation to the offspring are largely unknown. Here we show an example of intergenerational adaptive response that is induced in anticipation of a changing environment. The neuronal transduction of an environmental signal correlates with changes in chromatin modifications status in the parental germline, resulting in offspring with alternative phenotypes.

Phenotypic plasticity across generations is predicted to evolve when local environments can be anticipated, thus providing a means for the mother to adjust the phenotype of the offspring to enhance their success in that environment [22-24]. Although the ecology of *A*. *freiburgensis* is not known, a strain of this species (JU1782) has been isolated from rotting plant stems, which is similar to the *C*. *elegans* habitat [25]. In this environment, it is common a “boom-and-bust” type of lifestyle, characterized by rapid consumption of food sources and population growth, followed by a period of dispersal. Chemosensation of the environment by the nematode *A*. *freiburgensis* determines the developmental trajectory and sex of the F1 generation. Since *A*. *freiburgensis* dauers are migratory and develop into selfing hermaphrodites, colonizer individuals assure propagation in the new habitats even in the absence of conspecific males.

The chemosensory ASH neurons do not connect directly to the germline. Thus, it is likely that the endogenous signaling from sensory neurons to the germline is mediated by some neuroendocrine messenger that directly or indirectly affect the epigenetic status of the mitotic germline. The exposure of hermaphrodites to high-density conditions leads to higher levels of H3K4me3 and lower levels of H3K9me3, and correlates with higher transcription rates [15]. It remains to be determined which genes are activated and how they affect the change in sex determination of the offspring. Changes in the transcription of germline genes in response to environmental stimuli has been recently demonstrated in mammals as well [26, 27].

Although there is considerable variation in the neuroanatomy of nematodes [28], *Auanema* and *Caenorhabditis* are morphologically similar and phylogenetically close enough [6, 29] to make the identification of amphid neuron homologs relatively easily. In *C*. *elegans* and other nematodes of the Eurhabditis clade, the ASH neuron is a nociceptor that mediates avoidance behavior to harmful stimuli [14, 30, 31] and has been recently implicated in dauer induction [32]. Differences in dauer formation between *C*. *elegans* and *A*. *freiburgensis* (within and across generations, respectively) could be due to the expression of particular neurotransmitters in homolog neurons [33, 34]. Future comparative studies with closely related species will reveal how mechanisms of intergenerational inheritance evolved.

## Acknowledgements

A.P.-d.S. acknowledges funding by the National Science Foundation (IOS1122095), BBSRC (BB/L019884/1) and Leverhulme Trust (RPG-2016-089). We thank Nick Burton for critically reading the manuscript.

## Author contributions

G.Z., V.K., P.R. and A.P.-d.S.. designed the study. G.Z, V.K., J.C., C.B. and B.H. conducted experiments that involved the production of conditioned medium, crosses, and sexing offspring. G.Z. and V.K. performed laser ablations and P.R. performed the immunocytochemistry experiments. G.Z., V.K., P.R. and A.P.-d.S. wrote the paper.

## Author Information

The authors declare no competing interests. Correspondence and requests for materials should be addressed to A.P.-d.S.

## Supplementary Materials

## Methods

### Strain and culture

Unless otherwise stated, we used *A*. *freiburgensis* strain SB372 throughout this study [6]. The strain JU1782 was isolated from rotting *Petasites* stems sampled in Ivry (Val-de-Marne), France, in September 2009 by Marie-Anne Félix. Nematodes were cultured at 20 °C on standard Nematode Growth Medium (NGM) [35] plates seeded with the *Escherichia coli* OP50-1 strain. Microbial contamination was prevented by adding 25 μg/mL nystatin and 50 μg/mL streptomycin to the NGM.

### Isolation of age-synchronized hermaphrodite adults

To obtain adult hermaphrodites of about the same age, we relied on the finding that every *A*. *freiburgensis* dauer develops into a hermaphrodite. When many young hermaphrodites were required, crowded culture plates were treated with 1% sodium dodecyl sulfate (SDS) [11]. After this treatment, only dauer larvae remain alive. The treatment consisted of first resuspending nematodes in 2 ml of water for each plate. They were then transferred to 15 ml tubes and sedimented by centrifugation at 3,500 rpm for 3 minutes. After discarding the supernatant, 10 ml of 1% SDS was added to each tube. Nematodes were incubated in SDS at room temperature for 30 minutes, after which they were sedimented by centrifugation at 3,500 rpm for 3 minutes. 10 ml of water was added to each tube, and nematodes were resuspended and centrifuged at 3,500 rpm for 3 minutes. Dauers were transferred to a freshly seeded 6 cm plate and left to crawl out of the carcasses and debris. For isolation of a small number of young hermaphrodites, dauers were isolated from an overcrowded plate. They can easily recognized by their nictation behavior, in which they stand on their tails and wave their bodies.

### Sexing of offspring

To determine the sex of the F1 from selfing hermaphrodites, the hermaphrodite mothers were selected by first isolating dauers, which in *A*. *freiburgensis* always develop into hermaphrodites. Each dauer was placed on a 6 cm seeded NGM plate and kept at 20 °C to develop into an adult. Eggs laid by the hermaphrodite mother were collected and placed onto 96-well plates and left to develop until adulthood. Hermaphrodites were distinguished by their ability to produce offspring in the absence of a mating partner. Females typically lay unfertilized oocytes, and males are identified by their blunt tails [6]. When reporting sex percentages, we considered only the XX offspring (hermaphrodites or females).

### Production of conditioned medium from high-density nematode cultures

Plates of *A*. *freiburgensis* on NGM (with *Escherichia coli* OP50-1) were grown for ca. 6 days (20 °C) until a high population density was reached (usually considered > 1,000 nematodes/cm^2^) and washed with M9 medium [35] (10 ml, supplemented with 25 μg/mL nystatin) into a 1,500 ml flask. This procedure was repeated with several plates over 3 weeks, until 1,000 ml of liquid culture was produced. The liquid culture was during that time incubated on a rotary shaker at 22 °C and 100 rpm. Contamination was prevented by adding 25 μg/ml nystatin to the medium. Subsequently, the culture was centrifuged to sediment nematodes and bacteria, and the conditioned medium was collected and freeze-dried.

### Assay with conditioned medium

To test if *A*. *freiburgensis* hermaphrodites respond to environmental signals to produce different types of offspring, they were cultured in isolation either in the presence or absence of conditioned medium (prepared as described above). First, young hermaphrodite mothers were isolated by treating nematode culture plates with 1% sodium dodecyl sulfate (SDS), as described above. Dauers were placed into 6 cm plates seeded with *E*. *coli* OP50-1 for 22-24 hours and left to develop until the L4 larval stage. In assays with conditioned medium, each L4 hermaphrodite was placed into a 6 cm plate containing freeze-dried conditioned medium powder (50 mg), dissolved in 200 μl *E*. *coli* OP50-1 medium. After overnight incubation at 20 °C, eggs were collected and washed with 200-300 μl of M9. Each egg was then moved to a single well of a 96-well microtiter plate, in which each well contained 100 μl of NGM seeded with *E*. *coli*. Sex of the offspring was scored as described in the previous section. Each assay with the conditioned medium was performed side by side with a control (L4 hermaphrodite cultured in the absence of conditioned medium).

In the experiment to test if mothers react sense crowding cues, L4 animals (N= 10) were placed on 6 cm seeded plates with or without conditioned medium and left to develop to adulthood and to lay eggs for 16 hours. A total of 149 and 199 F1 offspring were sexed in the absence and presence of conditioned medium, respectively. In the test for induction of dauers for more than one generation, each of the F1 (N= 3) hermaphrodites were placed on single plates and the brood produced overnight was sexed after they developed into adults (N= 141 F2s).

To address whether L1 larvae can respond to conditioned medium, dauers were placed on a 6 cm plate seeded with *E*. *coli* for 22-24 hours. Each resultant L4 hermaphrodite was transferred to one 6 cm seeded plate and left to lay eggs. The offspring (at L1 stage) were placed onto 6 cm plates containing conditioned medium (as described above) and left to develop until adulthood, when they were scored for their sexual identity.

### Determining the minimum density of nematodes to induce hermaphrodite offspring

Young hermaphrodite adults were placed in 6 cm NGM plate seeded with *E*. *coli* OP50-1 lawn (0.7 cm radius). After 6 hours at 20 °C, the adult mothers were removed from the plate. Eggs were allowed to develop and were sexed as above. This experiment was performed in triplicate and the number of mothers tested were 10,15, 20, 25 and 30.

### Effect of temperature and starvation

To test the effect of temperature stress (25 °C) and starvation on the mother to induce dauer and hermaphrodite development, dauers were picked from a 6 cm agar plate spotted with a ∼5 mm diameter *E*. *coli* OP50-1 lawn that was kept at 20 °C. At early L4 stage each worm was moved to a 6 cm NMG plate spotted with *E*. *coli* OP50-1 and incubated at 25 °C or in the absence of food. Laid eggs were moved to a single well of a 96-well plate and incubated at 20 °C to allow them to develop into adults. Sexes of the offspring were determined as above.

### DiI staining

The red fluorescent lipophilic cationic indocarbocyanine dye DiI (1,1“-dioctadecyl-3,3,3”,3”-tetramethylindocarbocyanine perchlorate) (Molecular Probes - stock dye in dimethyl formamide solution containing 2 mg/ml (2 mM), was used to identify the amphid neurons. Hermaphrodites at L4 stage were washed from plates twice using M9 buffer [35], resuspended in 1 mL of M9 containing 8 μl DiI stock solution (1:125 final dilution) and incubated on a slow shaker at room temperature for at least 3 hours in the dark. After the incubation, nematodes were washed twice with M9 buffer to remove residual dye. Nematodes were then moved to a fresh NGM plate and were left to crawl on the bacterial lawn for at least 1 hour to allow clearance of ingested dye from their digestive tracts.

### The identification and ablation of amphids

A drop of melted 3% agarose was placed on a slide and flattened into a pad. Two slides containing spacers were used as guides for flattening the agar, so the thickness of the agar pad was equal to that of the spacer layer. In order to immobilize nematodes for imaging and surgery, sodium azide (an inhibitor of mitochondrial respiration) at a final concentration of 0.1 M was used in the agar pad. Nematodes were placed onto a drop of anesthetic (sodium azide) and a coverslip was placed on the slide. Care was taken to avoid creating bubbles next to the nematodes, as air-water interfaces can interfere with imaging and laser surgery.

Amphid neurons were identified according to the position of the cells viewed under Nomarski optics (Zeiss Axio Observer.Z1), using the appropriate filters (DiI gives red fluorescence). For ablation a diode laser-pumped pulsed dye laser (Micropoint laser ablation system, Andor) was used. According to the supplier, this laser unit generates a laser beam with peak wavelength of 435 nm, pulse energy up to 50 μJ, peak power 12 kW (average power 750 μW), peak duration 3-5 ns, pulse repetition rate 0-15 Hz. To confirm that a damage had occurred, features such as difference in the morphology of the cell before and after ablation and the change of refractive index in the nucleus were taken into consideration. Loss of fluorescence of the ablated cell was not considered sufficient to assume that a damage was induced, since this can be merely due to a photobleaching mechanism.

To rescue nematodes, the coverslip was removed from the slide very gently and 200 μl of M9 were used to move the worm to a seeded (*E*. *coli* OP50-1) NGM plate (25 μg/mL nystatin, 50 μg/mL streptomycin). Since nematodes were very dehydrated at this point, they were allowed to recover for 2 or 3 hours before proceeding with subsequent biological assays.

For the biological assay, each ablated L4 hermaphrodite was moved to a 6 cm plate containing freeze-dried supernatant powder (50 mg) dissolved in 200 μl of an overnight culture of E. coli OP50-1 in LB medium supplemented with 50 μg/mL streptomycin. Non-hatched eggs that were laid during the night were collected and were washed with 200-300 μl of M9 buffer. Each egg was moved to a single well of a 96-well microtiter plate (100 μl NGM per well). Sex of the F1 offspring was identified by checking for F2 offspring to determine if hermaphrodite, dead oocytes to determine if female and tail morphology for male determination. The procedure for mock ablations was identical (sodium azide treatment, fluorescent illumination, time in slide), except that no neuron was ablated.

### Gonads Immunohistochemistry procedures

Hermaphrodites at L4 stage were placed on a 6 cm NGM plate containing 50 mg of conditioned medium. After 24-36 hours, gonads were dissected on a slide (Superfrost microscope slide, VWR) in M9 buffer (22 mM KH_2_PO_4_, 34 mM K_2_HPO_4_, 86 mM NaCl, 1 mM MgSO_4_) and dissected gonads were fixed and treated with PBST (PBS + 0.05% Tween-20) and PBST + 0.5% BSA as a previously described [36]. Mouse primary antibodies to H3K9me3 (gift from Dr. H Kimura from Tokyo Institute of Technology) and H3K4me3 (CMA304, from Millipore 05-1339-S) were applied at a 1:2,000 dilution in PBST and incubated at room temperature overnight. Slides were washed twice in PBST for 10 minutes and a secondary antibody (goat, anti-mouse Alexa 478, Invitrogen) was applied at a 1:100 dilution in PBST for 2 hours at room temperature. Slides were washed in PBST as above to remove the excess of secondary antibody and then one drop of Fluoroshield Mounting Medium with DAPI (from Abcam ab104139) was added on the immunostained samples. Controls were not exposed to conditioned medium before performing gonad dissection and immunostaining. Images were taken with a 60X objective in 2.40 μm z stack intervals (12 sections) with a Deltavision microscope (Olympus).

### Treatment with IOX-1

For treatment with 5-carboxy-8-hydroxyquinoline (IOX-1, abcam ab144394), young adults nematodes were transferred and incubated for 24-48 hr in NGM plates containing IOX-1. IOX-1, previously diluted in dimethyl sulfoxide (DMSO) at 1 μM final concentration, was added to seeded NGM plates. Afterwards, nematodes were dissected as indicated in immunostaining protocol. Controls contained no IOX-1, but DMSO. To check IOX-1 effects in offspring, a single young adult was transferred to an NGM plate in the presence of IOX-1. Embryos were collected in 96-well plates during four days and the sex was scored as described above.

---

**Figure S1.**
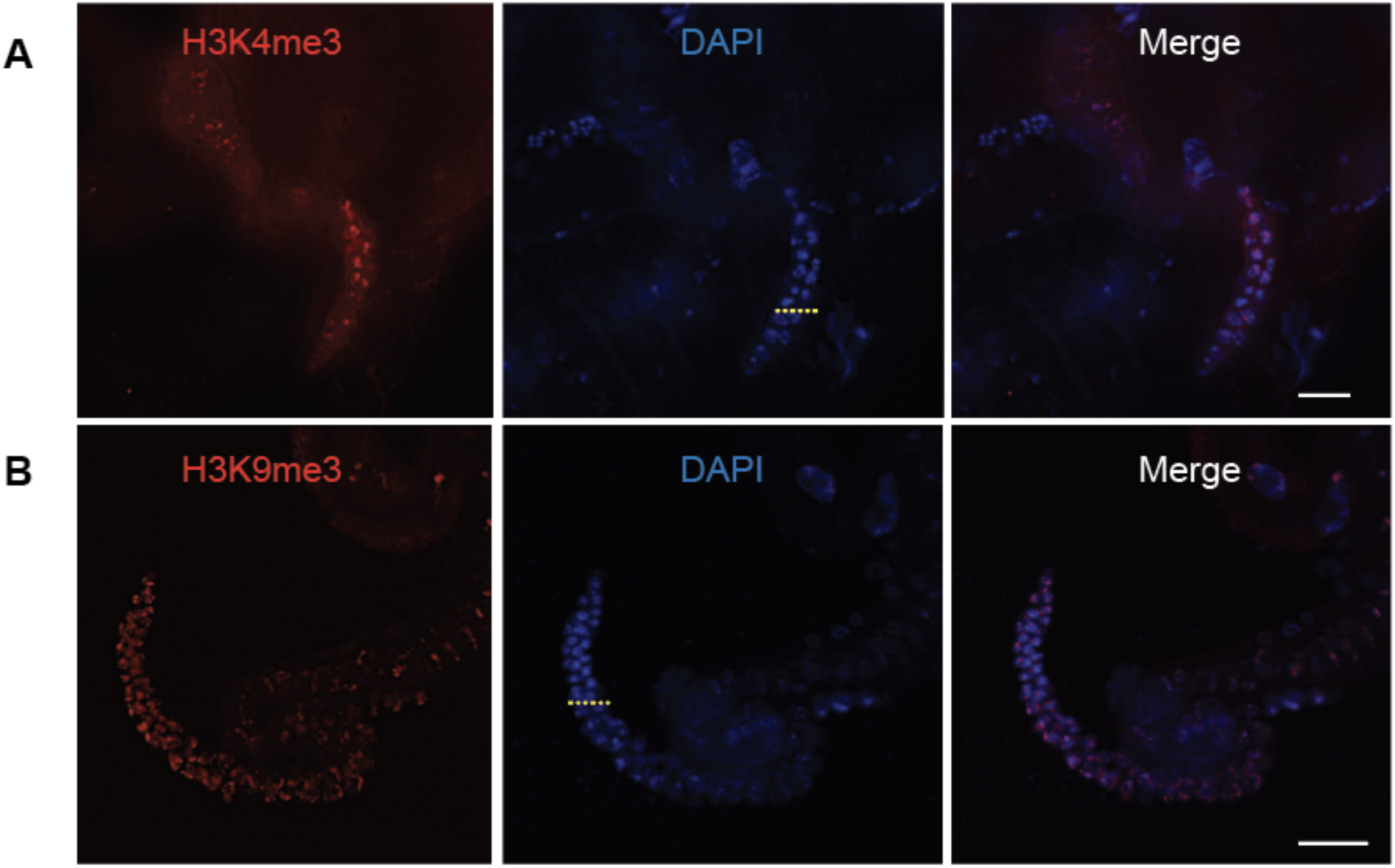
IOX-1 changes the histone methylation patterns in hermaphrodites in (+) CM plates. The mitotic germline has low levels of H3K4me3 (**A**) and high levels of H3K9me3 (**B**). The yellow dotted line in the pictures with DAPI staining represents the border between the mitotic and meiotic part of the germline. In (**A**) the mitotic part is to the bottom of the yellow line, and in (**B**) to top. Scale bar= 25 μm.

